# xAtlas: Scalable small variant calling across heterogeneous next-generation sequencing experiments

**DOI:** 10.1101/295071

**Authors:** Jesse Farek, Daniel Hughes, Adam Mansfield, Olga Krasheninina, Waleed Nasser, Fritz J Sedlazeck, Ziad Khan, Eric Venner, Ginger Metcalf, Eric Boerwinkle, Donna M Muzny, Richard A Gibbs, William Salerno

## Abstract

**Motivation:** The rapid development of next-generation sequencing (NGS) technologies has lowered the barriers to genomic data generation, resulting in millions of samples sequenced across diverse experimental designs. The growing volume and heterogeneity of these sequencing data complicate the further optimization of methods for identifying DNA variation, especially considering that curated highconfidence variant call sets commonly used to evaluate these methods are generally developed by reference to results from the analysis of comparatively small and homogeneous sample sets.

**Results:** We have developed xAtlas, an application for the identification of single nucleotide variants (SNV) and small insertions and deletions (indels) in NGS data. xAtlas is easily scalable and enables execution and retraining with rapid development cycles. Generation of variant calls in VCF or gVCF format from BAM or CRAM alignments is accomplished in less than one CPU-hour per 30× short-read human whole-genome. The retraining capabilities of xAtlas allow its core variant evaluation models to be optimized on new sample data and user-defined truth sets. Obtaining SNV and indels calls from xAtlas can be achieved more than 40 times faster than established methods while retaining the same accuracy.

**Availability:** Freely available under a BSD 3-clause license at https://github.com/jfarek/xatlas.

**Supplementary information:** Supplementary data are available at *Bioinformatics* online.

## Introduction

Improvements in the identification of single-nucleotide variation (SNVs) and small insertions and deletions (indels) from next-generation sequencing (NGS) data remains an active area of research. Small variant calling pipelines using different components and techniques have been able to achieve consistently high variant call accuracy rates, with surveys of variant calling methods finding accuracy rates exceeding 97% as of 2015 (Highnam *et al.*, 2015), and exceeding 99% as of 2016 (Altman *et al.*, 2016). Despite this, there are ongoing algorithmic developments to refine small variant calling to address new demands in research and clinical domains, such as the need for accurate and reproducible variant calls in clinical settings (Richards *et al.*, 2015), characterizing rare variants in common diseases (Cirulli and Goldstein, 2010), and identifying non-diploid variation (Campbell *et al.*, 2015).

At the same time, there are also growing disparities in size and heterogeneity between the data involved in current large-scale NGS experiments and the data used to build high-confidence (or “gold standard”) variant call sets that are widely used to evaluate variant calling methods. The Trans-Omics for Precision Medicine program (https://www.nhlbiwgs.org/), for example, has sequenced more than 54,000 whole genomes to date and aims to sequence at least 120,000 samples in total. The overall volume and heterogeneity of these data well exceeds those for both the highly curated individual samples used to create high-confidence variant call sets, such as those from the Genome in a Bottle project (Zook *et al.*, 2014), and even broader analyses of genetic variation that have been carried out by earlier large-scale studies, such as the 1000 Genomes Project (The 1000 Genomes Project Consortium, 2015). The focus of existing methods on managing the scale of current challenges has also been at the expense of considerations of the diversity of experimental designs that can affect variant call quality. Such factors include the analysis of DNA samples of various quality, heterogeneous tissue types, different library preparation and variable read lengths, insert sizes and read alignment, and analysis filtering parameters.

Considerable efforts have also been focused on increasing the computational efficiency of sequence analysis and variant calling at scale, with GATK (McKenna *et al.*, 2010), DeepVariant (Poplin *et al.*, 2017), Edico Genome’s Dragen (Miller *et al.*, 2015), and other state-of-the-art methods leveraging distributed software and hardware-optimized technologies. However, these methods often require commitments to external infrastructure or internal technology development to be applied effectively, which may not be well-suited for rapid turnaround times or cost-effective execution of variant calling at scale.

The ideal variant caller would therefore manage both scalability issues and sources of data heterogeneity. Further, the method should also allow rapid turnaround times for the generation of single-sample call sets so that different parameters can be evaluated. We have previously reported the software packages ATLAS (Havlak *et al.*, 2004) for DNA assembly and Atlas2-SNP (Shen *et al.*, 2010) and Atlas2-INDEL (Challis *et al.*, 2012) for small variant calling. Here, we describe xAtlas, an accurate and fully-featured variant calling application that requires only a fraction of the computational resources of other small variant calling methods.

## 2 Methods

xAtlas is a command-line C++ application that employs HTSlib (Li *et al.*, 2009) to handle alignment and variant call file formats. Input sample alignment files may be in either BAM (Li *et al.*, 2009) or CRAM (Fritz *et al.*, 2011) format. When writing output in VCF format (Danecek *et al.*, 2011), xAtlas may optionally include non-variant VCF entries spanning regions not covered by variants formatted in genome VCF (gVCF) format (https://sites.google.com/site/gvcftools/) to facilitate downstream multi-sample variant analyses. The application also may be built with multithreading support, which allows the processes of reading the input alignment file, processing SNVs, and processing indels to be handled each in a separate thread.

### 2.1 Variant Detection and Evaluation

xAtlas variant calling is performed in the following high-level stages: preliminary read filtering; collecting candidates for variant calls from the alignment file; evaluating each candidate variant; and reporting candidate variants (Figure 1). As reads are scanned from the input alignment, a read will be filtered out from variant calling if it is marked as an unmapped, as a duplicate read, or alternatively has a mapping quality score below a minimum threshold, with a default of one. Candidate sequence variations are then collected from the unfiltered reads and grouped for variant calls. To aggregate candidates, sequence variations are identified within each read by locating the coordinates at which sequences differ from the provided reference genome. The SAM format’s CIGAR string, which defines the edit operations between the read’s sequence and the reference sequence at its mapped position, is used to determine variant coordinates. SNVs are defined as point differences between reference and aligned sample sequences within the spans of CIGAR match operators. Variant alleles are assigned reference coordinates that correspond to its parsimonious representation within the alignment, as defined by Tan *et al.* (2015).

**Fig. 1.**
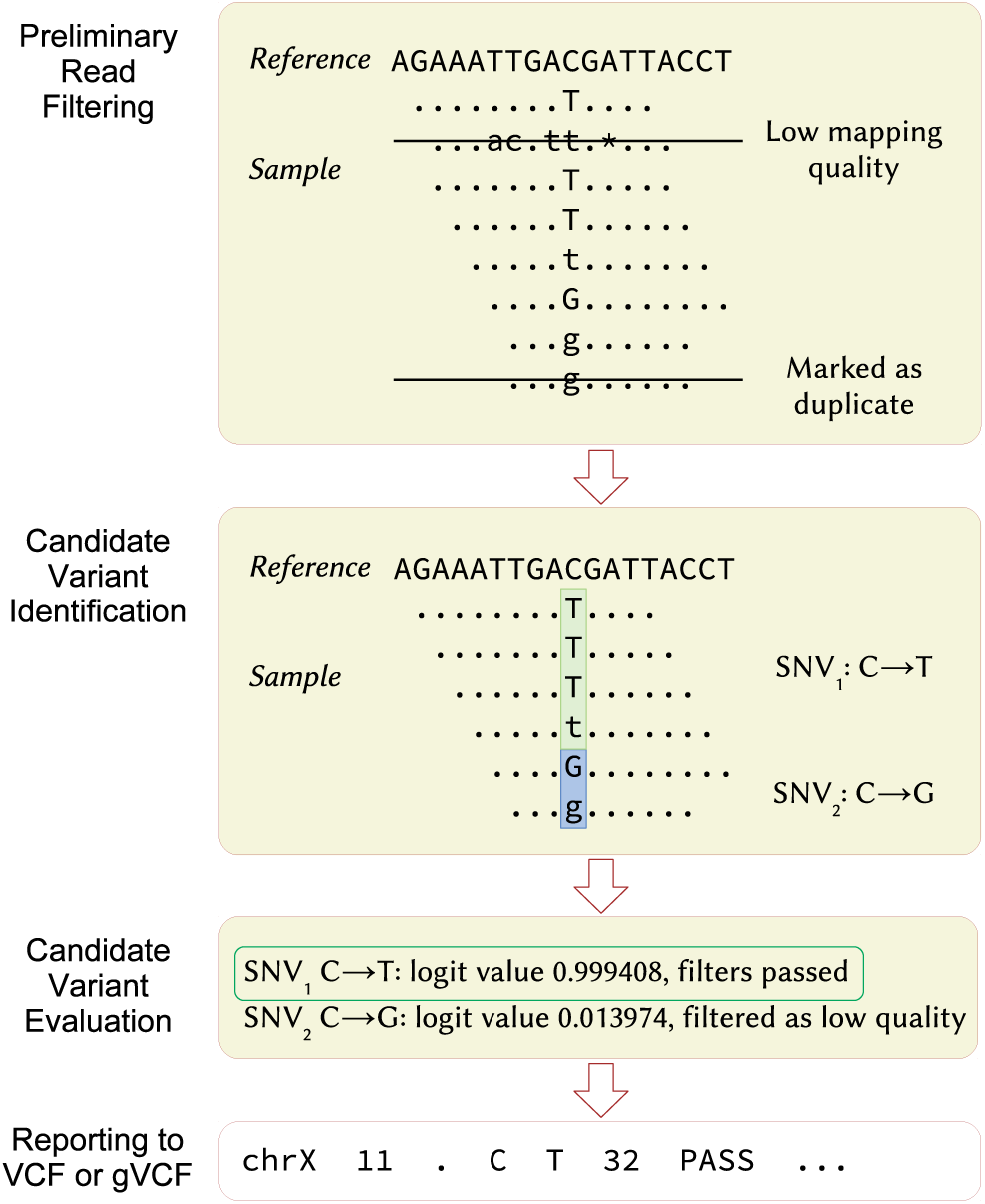
Variant calling in xAtlas is performed in three stages: filtering out undesirable reads, identifying and grouping alleles into candidate variants, and evaluating the likelihood of each candidate variant.

While collecting candidate variants to evaluate, xAtlas also records multiple sequence and alignment attributes associated with each candidate variant, including base quality values and coverage ratios for reads supporting the reference sequence vs. those supporting the variant allele. These features are evaluated together to assign the a confidence score for the candidate variant, utilizing either a SNV or indel candidate variant evaluation model based on logistic regression. xAtlas allows the user to redefine the logistic regression intercepts and variable coefficients of these models with values that may be derived from retraining on new samples. Thresholds for both confidence scores and candidate variant features determine whether a candidate variant will be called and if a called variant will be filtered in the output VCF file. Other filters may also applied to candidate variants based on other features not evaluated by the candidate evaluation models, such as if there are too few reads supporting the candidate variant allele.

After assigning variant confidence scores from the logistic regression models and applying filters, xAtlas then determines the most likely then genotype and reports the candidate variant in the VCF. A variant call is reported in the VCF only if the candidate’s logistic regression value is greater than an adjustable cutoff, with a default value of 0.25. If multiple variants may be reported at the same position, xAtlas reports only the variant at that position with the greatest number of reads supporting the variant sequence. For SNVs, if there are still multiple candidates tied for the greatest number of supporting reads, the candidate variant with the highest logistic regression value is then selected. xAtlas assigns the genotype 1/1, 0/1, or 0/0 to called variants. For indel, genotypes are assigned based on cutoffs for the ratio of reads supporting the variant allele to the total number of reads overlapping the indel. For SNVs, each SNV is assigned the genotype with the highest genotype likelihood as determined by xAtlas (Supplementary Information Section 1).

### 2.2 Retraining Candidate Variant Evaluation Models

The logistic regression model retraining performed as part of this study was performed by building sets of positive and negative examples of variant sites from pairs of sample alignments and using subsets of these variant sites in logistic regression model fitting. The set of all possible candidate variant sites and the values that xAtlas supplies to the SNV and indel logistic regression models were compiled for each sample. Subsets of positive and negative variant site examples were then derived from this set based on variant site overlap with a truth set of high-confidence variants and with high-confidence variant regions. Positive variant sites were selected from variant sites present in both technical replicates, overlapping the NIST high-confidence variants, and restricted to the NIST high-confidence regions. Two types of negative variant sample sites were compiled, where variant sites are either present in both technical replicates or present in only one of the two replicates, with both types restricted to the NIST high-confidence regions but not overlapping the NIST high-confidence variants. Each of these comprised half of the negative example variant sites in assembled training and testing sets. Training and testing sets were compiled as non-overlapping sets of 10,000 randomly sampled positive and negative variant site examples, with a 1:1 ratio of positive vs. negative examples in each set. Logistic regression model fitting using these training and testing sets was performed using the LogisticRegression classifier from scikit-learn (Pedregosa *et al.*, 2011).

## 3 Results

### 3.1 Variant Call Quality Assessment

xAtlas variant call quality was assessed by comparing variants called on alignments for multiple samples with multiple retrained logistic regression model parameters to NIST high-confidence variant sets. Seventeen sets of NA12878 WGS technical replicates or pairs of alignments sequenced on the Illumina HiSeq X and Illumina NovaSeq platforms, which include down-sampled alignments, were used to generate retrained logistic regression model parameters (Supplementary Table S1). These samples and additional samples of NA12878 and HG002 were used to evaluate the sets of trained logistic regression parameters. All samples sequenced at the Baylor College of Medicine Human Genome Sequencing Center (BCM-HGSC) were aligned with BWA-MEM (Li and Durbin, 2009) version 0.7.12 and had indels realigned using GATK IndelRealigner (McKenna *et al.*, 2010) version 3.4.0. For all sample sets, version 3.3.2 of either the NA12878 or HG002 high-confidence variants using GRCh37 reference coordinates from the NIST Genome in a Bottle Consortium (Zook *et al.*, 2014) and the accompanying high-confidence variant regions were used as the truth set to refine variant sites for positive and negative examples during retraining.

Sensitivity was calculated as the rate of concordance with the NIST high-confidence set, as measured by RTG vcfeval version 3.8.4 (Cleary *et al.*, 2015). Variants from the NIST high-confidence set were restricted for this comparison to those within high-confidence regions and to variants which were listed as not having multiple variant alleles or sequences associated with the same parsimonious position in the VCF. Figure 2 shows plots of precision against sensitivity for SNVs and indels. For the sample with the highest sensitivity and precision values, a NA12878 WGS sample sequenced on an Illumina NovaSeq sequencer with an average depth of coverage of 54.5, sensitivity was measured at 94.82% for indels and 99.85% for SNVs, and precision was measured at 91.19% for indels and 99.63% for SNVs. F-score measures of variant accuracy for published precisionFDA Truth Challenge (https://precision.fda.gov/challenges/truth) entries range from 97.77% to 99.96% for SNVs whereas xAtlas achieved 98.70% to 99.74% for SNVs. For indels, precisionFDA reported F-scores from 70.50% to 99.40% while xAtlas ranged from 84.76% to 92.97%, depending on coverage. With HG002, xAtlas achieved an F-score accuracy of 99.57% for SNVs and 93.07% for indels.

**Fig. 2.**
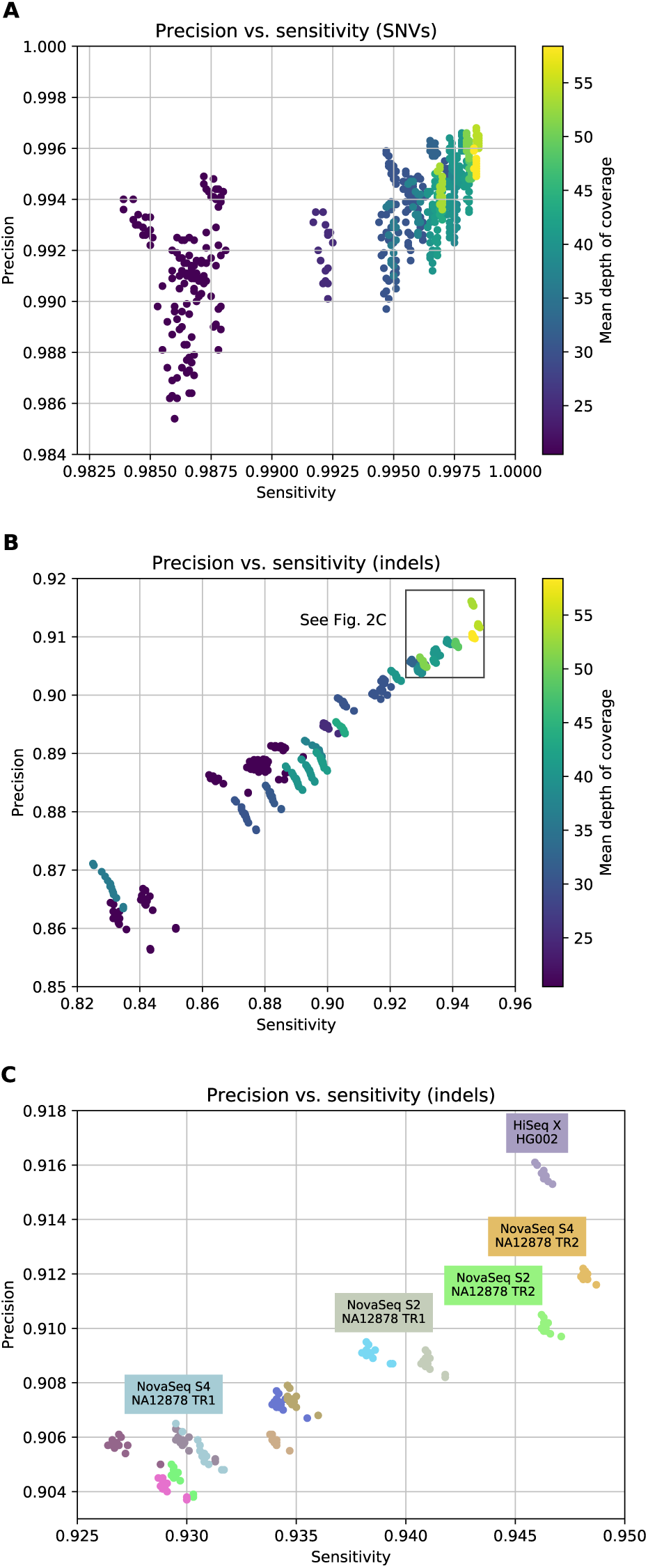
Sensitivity vs. precision for xAtlas SNVs (A) and indels (B) compared to the NIST NA12878 high-confidence variant set v3.3.2. Each point represents the sensitivity and precision of variants called by xAtlas using one of the retrained set of logistic regression parameters for its variant evaluation models. Clusters of similar sensitivity and precision values become apparent when these data points are grouped by sample (C).

### 3.2 Performance Benchmarking

Table 1 shows runtime and memory benchmarks of xAtlas runs with samples of varying coverage levels and with various runtime options. All runtimes were measured using a compute node with an Intel Xeon processor (Haswell series) in a high-performance computing cluster at the BCM-HGSC. When run as a single-threaded operation, xAtlas produced SNV and indel VCFs for an Illumina HiSeq X WGS sample at 30× coverage with a walltime of 2,096 seconds. Total memory usage remained stable across all samples and coverage levels, with resident memory usage averaging 3.7 GB and not exceeding 4 GB across all runs. For comparison we used a best-practices variant calling pipeline based on Samtools and BCFtools that generated output in VCF format. Runtimes for this Samtools and BCFtools variant calling pipeline across alignments ranging from approximately 20× to 50× coverage took on average 41 times longer (with an approximately 24 hour difference for the sample with 30× coverage) to run than xAtlas on the same sample, with runtimes for this pipeline also listed in Table 1.

**Table 1.** Runtimes for xAtlas and a Samtools/BCFtools-based small variant calling pipeline in HH:MM:SS and fold decrease in runtime from the Samtools/BCFtools-based pipeline to xAtlas when generating output in VCF format.

| Sequencing platform | Sample type | Total reads | Whole-genome coverage | Samtools/BCFtools | xAtlas (gVCF) | xAtlas (VCF) | Fold decrease (VCF) |
| --- | --- | --- | --- | --- | --- | --- | --- |
| Illumina HiSeq 2500 | Panel | 4,399,370 | N/A | 0:06:57 | 0:02:45 | 0:01:00 | 6.95 |
| Illumina HiSeq 2500 | Whole exome | 119,003,254 | N/A | 2:52:22 | 0:16:13 | 0:11:45 | 14.67 |
| Illumina HiSeq X | Whole genome | 452,321,930 | 20.51 | 18:24:38 | 0:31:58 | 0:25:12 | 43.83 |
| Illumina HiSeq X | Whole genome | 673,989,846 | 30.51 | 24:33:56 | 0:42:38 | 0:34:52 | 42.27 |
| Illumina HiSeq X | Whole genome | 911,872,368 | 41.35 | 30:54:37 | 0:55:39 | 0:49:39 | 37.35 |
| Illumina NovaSeq | Whole genome | 1,045,849,198 | 51.18 | 43:50:00 | 1:06:05 | 1:03:59 | 41.10 |

Read lengths were 101 bp for Illumina HiSeq 2500 and 151 bp for Illumina HiSeq X and Illumina NovaSeq. Capture design sizes totaled 534,703 bp for panel and 35,810,763 bp for whole exome.

### 3.3 Applications of xAtlas in Large-scale Variant Analyses

To highlight the versatility of xAtlas as a method to identify SNV and indel calls across multiple scenarios, we describe three real use cases where we applied xAtlas at the BCM-HGSC.

First, xAtlas was was run on a whole-exome data set comprising of 15,000 BAMs (approximately 10 GB/sample) that is routinely used to annotate other samples. xAtlas allowed us to create gVCFs for these samples in 6708 total core hours, resulting in a single 1.5 TB resource for cost-efficiently harmonizing this set with new samples as needed.

Second, we performed a similar analysis on 22,609 whole-genome CRAMs (approximately 25 GB/sample) in parallel with multi-method structural variant analysis. Its low memory and CPU usage allowed us to run xAtlas for no additional cost by incorporating it in an existing cloud-based structural variant application that executes multiple structural variant and quality control methods simultaneously on a 32-core AWS instance.

Third, we used xAtlas to assess putative mosaic sites in 16,000 clinical panel BAMs (approximately 6 GB/sample). We modified xAtlas runtime parameters to report all evidence of variation in gVCF format, similarly to a traditional pileup. These gVCFs were then merged into a project-level VCF and filtered for putative mosaic sites by examining multi-sample profiles. Creating all 16,000 gVCFs took less than 2 walltime hours when split naively across 500 4-core AWS instances, allowing for inexpensive and rapid adjustment of variant calling parameters based on iterative assessment of the full data set without necessarily having to optimize for cloud instance types or minimizing cloud-specific data-transfer overheads.

## 4 Discussion

The development of xAtlas has produced an accurate, scalable, and retrainable DNA variant calling method. While its variant calling behavior has been shown to be both accurate and adaptable through model retraining, its utility as a component in a variant calling pipeline derives immensely from its computational efficiency. With runtimes of one CPU-hour per 30× whole-genome short read alignment, xAtlas has permitted variant analysis at the BCM-HGSC to scale to sample sizes that would have previously posed much greater challenges in terms of both computational resources and analytical interpretation.

Different alignment properties were found to variously affect the measured rates of sensitivity and precision of variants called by xAtlas. Across all variant call sets produced from all samples, xAtlas variant precision is highly positively correlated with sensitivity, with a Pearson correlation of *r* = 0.97. Precision and sensitivity rates are more closely clustered by sample (within each coverage level) than by which specific retrained logistic regression parameter set was used, suggesting that sample-specific alignment characteristics are a greater determining factor of variant call quality than variant evaluation model retraining. The use of retrained candidate evaluation model parameters produced measurable, but less pronounced, differences in rates of variant sensitivity and precision within different runs on the same sample. Depth of coverage also has a clear effect on sensitivity and precision rates. Differences in variant sensitivity and precision within the same sample were more pronounced in samples with lower coverage levels. Variant sensitivity and precision rates steadily improve across all samples as coverage level increases, with the highest sensitivity and precision values recorded for samples with coverage levels of at least 50×.

xAtlas is most similar in functionality to variant calling methods that have similar hardware requirements and entry points (i.e. a “finished” BAM or CRAM alignment). As such, xAtlas does not implement read-based refinements such as indel realignment or base quality score recalibration. The customizable and rapid generation of gVCFs allows for development at scale, which is critical when analyzing data sets larger than those defined for established best practices. While xAtlas can be used as a supplement or replacement for other variant calling pipeline steps, one important feature that has not yet been fully implemented in xAtlas is the ability to report multiple variant alleles for a single variant call. Since the variant calling processes for SNVs and indels in xAtlas are not complementary (indels are not recognized by the SNV calling procedure, and *vice versa*) accurate variant discovery and evaluation for multi-allelic variant calling would require SNVs and indels to be processed concurrently. Modifying xAtlas’ candidate variant evaluation model to support multiple variant alleles should not only produce more accurate genotypes by allowing accurate genotyping of multiple variant alleles at a given site, but also allow more comprehensive retraining and variant evaluation.

Since xAtlas variant call quality was evaluated using largely NA12878 datasets (primarily NA12878 WGS samples sequenced on the Illumina NovaSeq, HiSeq X, and HiSeq 2500 platforms, and comparing to the NIST GIAB for NA12878), expanding retraining and evaluation to include a more diverse set of samples, sequencing platforms, and application types should allow a more complete assessment for how these factors affect the efficacy of retraining the logistic regression models and possibly which sequence or alignment features have the greatest effects on variant call quality.

## 5 Conclusion

xAtlas has demonstrated a combination of computational efficiency and variant call accuracy. Sensitivity and precision rates for both SNVs and indels called by xAtlas rank among those of other variant calling methods that have been used in practice. xAtlas has permitted fast and cost-effective variant analysis across multiple projects at the BCM-HGSC consisting of tens of thousands of whole-genome samples. For small or large-scale variant analysis, xAtlas can be scaled run in compute environments ranging from a single laptop to large HPC clusters or arrays of cloud instances. With the ability to generate VCFs and gVCF-formatted variant call sets in terms of minutes or hours per sample, development of new variant analysis methods can also be carried out with rapid turnaround rates.

